# Glycosomal phosphoenolpyruvate carboxykinase CRISPR/Cas9-deletion and its role in *Trypanosoma cruzi* metacyclogenesis and infectivity in mammalian host

**DOI:** 10.1101/2024.12.20.629751

**Authors:** Carolina Silva Dias Vieira, Wei Wang, Fernando Sanchez-Valdez, Jihyun Lim, Brooke E White, Camilla Garcia da Silva Souza, Rick L Tarleton, Marcia Cristina Paes, Natalia Pereira Nogueira

## Abstract

*Trypanosoma cruzi*, the causative agent of Chagas disease, possesses glycosomes - unique organelles that house key metabolic enzymes, several of which are promising therapeutic targets. Among them, phosphoenolpyruvate carboxykinase (PEPCK) plays a central role in succinic fermentation, the main pathway for NAD^+^ regeneration within the organelle. Using CRISPR/Cas9 editing, PEPCK gene was disrupted in *T. cruzi*, producing single-allele knockout epimastigotes (TcPEPCK-sKO) with reduced enzyme activity. This disruption impaired glucose consumption and mitochondrial respiration, particularly oxidative phosphorylation, reducing dependence on mitochondrial ATP production. To compensate, pyruvate phosphate dikinase was upregulated, increasing alanine production, possibly to maintain redox balance. Although TcPEPCK-sKO epimastigotes exhibited a minor reduction in growth, their differentiation (metacyclogenesis) and invasion were severely compromised. However, once inside the host cell, TcPEPCK-sKO amastigotes increased their replication, leading to enhanced trypomastigote production. The same was observed in *in vivo* infection, where TcPEPCK-sKO infection in IFNγ-deficient mice caused uncontrolled parasitemia and severe pathology, highlighting the PEPCK critical role in host-pathogen interactions. However, an intact immune system effectively contained TcPEPCK-sKO infection. Taken together, our findings demonstrate that PEPCK is crucial for *T. cruzi* energy metabolism, enabling the parasite differentiation within the insect vector and controlling the infection of mammalian host cells.

## Introduction

*Trypanosoma cruzi*, the protozoan causative agent of Chagas disease (CD), presents significant challenges to global health owing to its debilitating effects and high lethality. Despite more than a century since its discovery in 1909 **[1]**, CD remains a neglected disease, with an estimated eight million people infected worldwide. Each year, migration and alternative transmission routes contribute to approximately 30,000 new cases, resulting in more than 14,000 deaths. It is the most prevalent parasitic disease in the Americas, placing 75 million people at risk **[2-5]**.

Historically, the genetic complexity of *T. cruzi* has hindered the development of effective therapies and a preventive vaccine, as this protozoan genomic intricacy precludes the use of traditional molecular genetic tools. This has left many of its biological processes obscure and it is significantly understudied compared to its relative, *Trypanosoma brucei* **[6]**.

The advent of CRISPR/Cas9 technology has revolutionized this field **[7]**, offering a potent and precise method for gene editing that was previously unattainable. This technology not only facilitates the efficient suppression of multiple genes but also opens new avenues for probing the intricate biology of this pathogen. It has been proven to be instrumental in dissecting the functional roles of various proteins and metabolic pathways, validating antiparasitic targets, and deepening our understanding of the pathogenic mechanisms of trypanosomatids **[8-11]**.

In terms of energy metabolism, *T. cruzi* is capable of metabolizing carbohydrates, amino acids, lipids, and glycerol, depending on the conditions to which it is subjected, and excretes different molecules resulting from the breakdown of these nutrients (succinate, acetate, and alanine) **[12-16]**. Glucose consumption is associated with the unusual fact that even under aerobic conditions, these protozoans partially oxidize glucose and excrete still- reduced compounds, mostly succinate, into the environment. This metabolic route is present in glycosomes, which are organelles exclusive to trypanosomatids, and is called succinic fermentation **[12, 17]**. Glucose is transported to glycosomes when taken up from the extracellular medium, where glycolysis begins in trypanosomatids. In this organelle, glucose is converted to 1,3-bisphosphoglycerate, which is then transferred to the cytosol, where it undergoes a series of reactions until the generation of phosphoenolpyruvate (PEP). For succinic fermentation to occur, PEP is transported from the cytosol to glycosomes and, after catalysis by phosphoenolpyruvate carboxykinase (PEPCK) and subsequent redox reactions, generates succinate, which is finally secreted into the environment.

As glycosomal enzymes have unique functional and structural characteristics that differ from those of mammals, they are attractive targets for the discovery of antiparasitic drugs. For this purpose, we selected and deleted the genes involved in glucose metabolism and succinic fermentation, PEPCK, through CRISPR/Cas9 and meticulously studied the functional and phenotypic effects of this protein knockout in this parasite to broaden the understanding of the biology and pathogenesis of *T. cruzi*.

## Results

### Generation of knock out for PEPCK gene using CRISPR/Cas9 system

In glycosomes, PEPCK catalyzes the carboxylation of phosphoenolpyruvate (PEP) to oxaloacetate to generate ATP. This is the first succinic fermentation reaction and is the main pathway responsible for regenerating NAD^+^ inside the organelle, which is reduced during glycolysis **[12]** (Figure 1A). To investigate the role of PEPCK in the *T. cruzi* life cycle, deletion of this gene in epimastigotes was induced using the CRISPR/Cas9 system. The parasites were transfected with ribonucleoprotein complexes (RNP) with *in vitro* transcribed sgRNAs targeting all gene copies in the genome. A repair template containing three in-frame stop codon sequences and a short sequence of M13 bacteriophages flanked by short homology regions was inserted into the cut site, promoted by the endonuclease (Figure 1B). The knockout of PEPCK was validated by PCR using primers for the PEPCK ORF and M13 bacteriophage sequence inserted after gene editing (primer sequences are shown in Table S1). The editing only generated single-allele mutant epimastigotes (Figure 1C). In accordance with the partial removal of the PEPCK gene (TcPEPCK-sKO), enzyme activity was reduced by 49% in epimastigotes (Figure 1D). In this study, the enzymatic reaction measured was the enzyme-mediated carboxylation of PEP into oxaloacetate, which produces metabolites in the succinic branch (Figure 1A).

**Figure 1.**
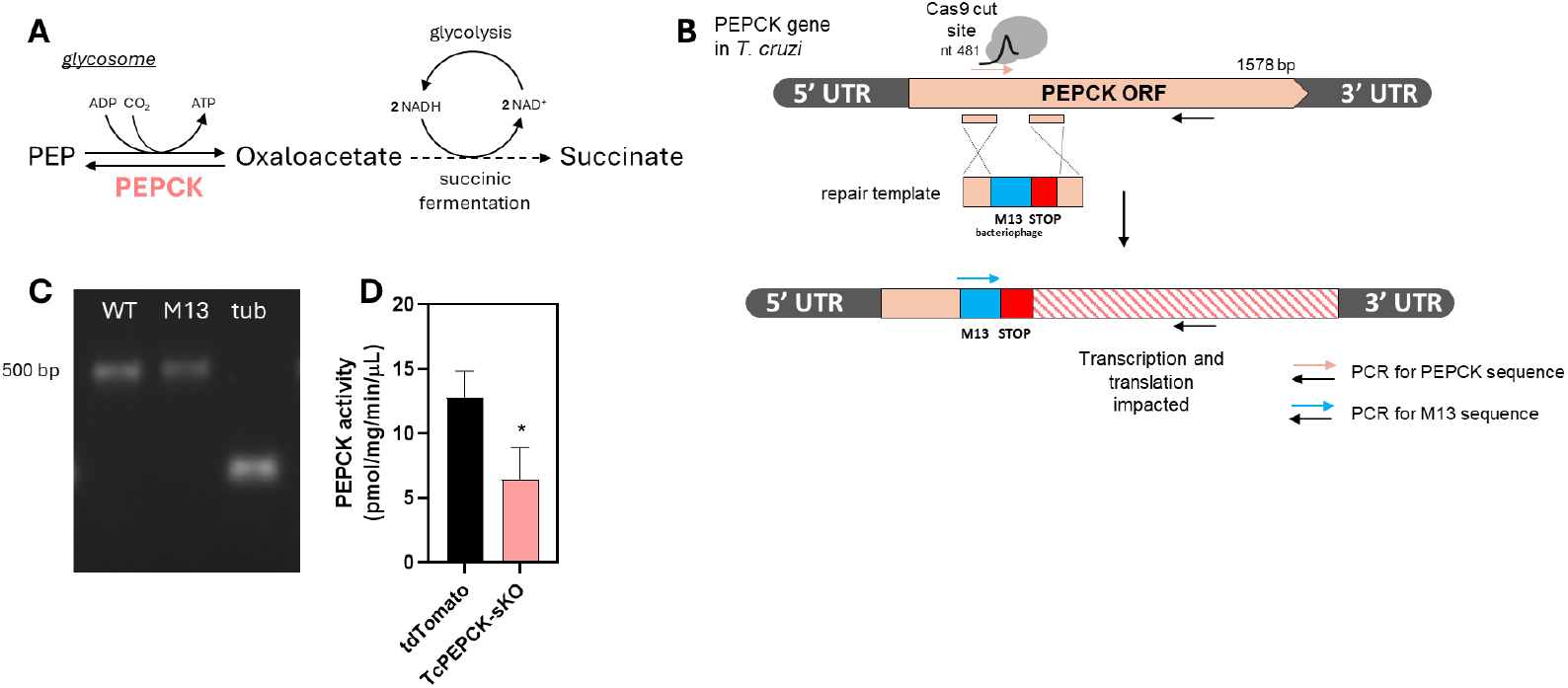
Strategy to generate PEPCK knockouts in *T. cruzi*. **(A)** Glycosomal PEPCK reaction in *T. cruzi*: Inside the glycosome, PEPCK is responsible for the first reaction of the succinic fermentation pathway. The enzyme catalyzes the carboxylation of PEP in oxaloacetate (OAA), generating ATP. OAA is then reduced in two subsequent reactions of the succinic fermentation route (glycosomal malate dehydrogenase-fumarate hydratase- fumarate reductase; *dashed line*) promoting the reoxidation of the glycosomal NAD^+^ and generating the final product, succinate. **(B)** Schematic representation of the strategy used to generate PEPCK knockout parasites mediated by CRISPR/Cas9. Cas9 induced a double-strand break (DSB) in the DNA at nucleotide 481 of the PEPCK open read frame (ORF). A repair template containing the M13 sequence (blue) and 3-in frame STOP sequence (red) flanked with homology sequences (from the PEPCK ORF) was supplied for the homologous recombination. **(C)** Validation of the PEPCK gene deletion by PCR. WT: primer sequence design for the cut site in the homology region; M13: specific primer for M13 bacteriophage inserted in the genome after the DSB-mediated by Cas9; Tub: tubulin DNA load control. **(D)** PEPCK activity quantified in tdTomato and TcPEPCK-sKO epimastigotes. The result shows mean ± SEM of five independent experiments. *p<0.05 related to tdTomato epimastigotes analyzed by Student t-test.

### The impairment of PEPCK gene affected *T. cruzi* epimastigotes bioenergetics

The growth of tdTomato and TcPEPCK-sKO epimastigotes was also evaluated in liver infusion tryptose (LIT) medium supplemented with or without 10 mM glucose. After seven days, no change was observed in the proliferation of tdTomato and TcPEPCK-sKO parasites maintained in glucose-depleted medium. When 10 mM glucose was added to the medium, the proliferation of both parasites increased proportionally, although TcPEPCK- sKO epimastigotes showed a 17% reduction compared with tdTomato parasites (Figure 2A).

**Figure 2.**
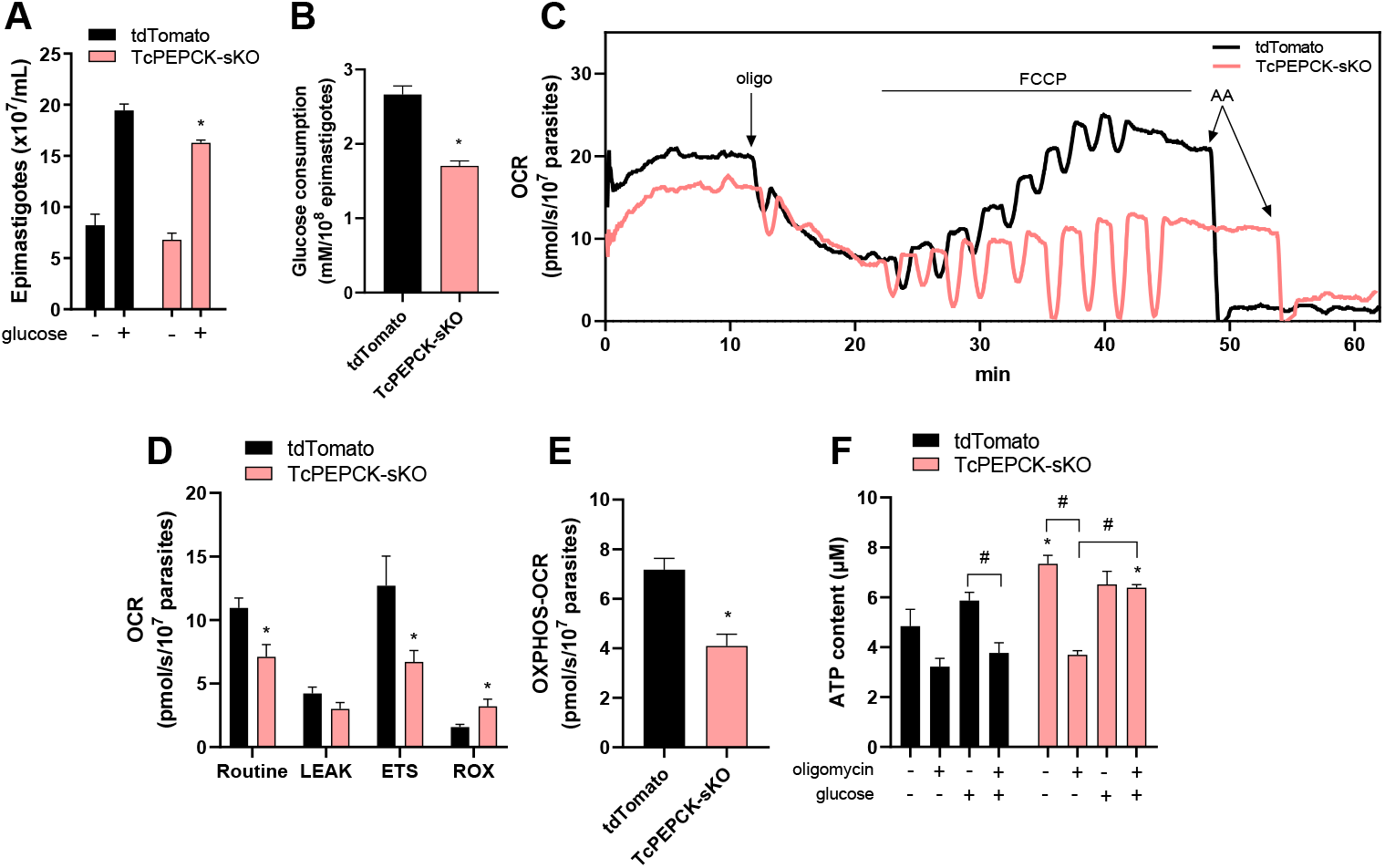
Proliferation, glucose consumption, and bioenergetic changes of TcPEPCK- sKO epimastigotes. **(A)** tdTomato and TcPEPCK-sKO epimastigotes were cultivated in LIT medium at 28 ºC, with or without the supplementation of 10 mM glucose for 7 days. The parasite proliferation was quantified using a Neubauer chamber. Representative data present mean ± SD of three independent experiments prepared in duplicates. *p<0.05, compared to tdTomato, analyzed using Student’s t-test. **(B)** tdTomato and TcPEPCK-sKO epimastigotes were incubated in LIT medium for 24 h. The glucose concentration of the supernatant was quantified using the Glucose Liquiform kit and normalized to the parasite number. Data are presented as the mean ± SEM of three independent experiments. *p<0.05, compared to tdTomato, analyzed using Student’s t-test. **(C)** The oxygen consumption rate (OCR) of tdTomato and TcPEPCK-sKO epimastigotes was evaluated using high-resolution respirometry. Representative OCR traces of epimastigotes indicating the addition of oligomycin (oligo), titration with FCCP, and antimycin A (AA). **(D)** Routine OCR, proton leak after oligomycin addition, maximal oxygen consumption (electron transport system-related OCR; ETS) after FCCP titration, and residual respiration (ROX) obtained after AA. **(E)** OXPHOS-related oxygen consumption related to ATP production (OCR) was calculated as the difference between routine and proton leak. The data are the mean ± SEM of four independent experiments. *p<0.05, compared to tdTomato, analyzed using Student’s t-test. **(F)** tdTomato and TcPEPCK-sKO epimastigotes were maintained in LIT medium without glucose supplementation for 24 h. Then, the parasites were incubated with or without 10 mM glucose and 2 μg/mL oligomycin for 24 h. After incubation, the intracellular ATP levels were evaluated using the ATPlite Luminescence ATP Detection Assay System. The data are the mean ± SEM of three independent experiments performed in duplicates. *p<0.05, compared to tdTomato, analyzed using Student’s t-test. #p<0.05, compared to each treatment, analyzed by One-Way ANOVA with post-hoc Tukey’s test.

Energy metabolism of tdTomato and TcPEPCK-sKO epimastigotes were also evaluated. Glucose consumption by these parasites was measured after 24 h of incubation in LIT medium. The results showed that the impairment of PEPCK promoted a decrease in 36% of the glucose consumption by the TcPEPCK-sKO parasite compared to tdTomato (Figure 2B).

Once the glucose consumption was compromised, mitochondrial respiration was assessed using high-resolution respirometry. TcPEPCK-sKO parasite showed a reduction in the Routine O_2_ consumption rate (OCR) compared with tdTomato mice (Figure 2C, 2D). Oligomycin was added to inhibit respiration related to ATP production (oligomycin- sensitive respiration). This inhibitor blocks ATP production from ATP synthase in the mitochondria, and the remaining OCR is from proton leak (oligomycin-insensitive respiration) (Figure 2C). Although proton leak was similar in both parasites (Figure 2D), OXPHOS respiration, which is related to mitochondrial ATP synthesis, decreased by 42% in TcPEPCK-sKO parasite (Figure 2E).

The maximal respiratory rate (OCR-related to the electron transfer system, ETS) was stimulated by adding increasing concentrations of FCCP, which uncoupled oxidative phosphorylation from ETS (Figure 2C). TcPEPCK-sKO epimastigotes also exhibited a 47% reduction in maximal O_2_ consumption (Figure 2D). Residual oxygen consumption (ROX) represents the non-mitochondrial respiration that is achieved after the addition of a complex III inhibitor, Antimycin A (AA). ROX increased 2-fold in TcPEPCK-sKO parasites compared to tdTomato (Figure 2C, 2D).

We also evaluated ATP production in the parasites challenged with glucose and oligomycin (Figure 2F). Without glucose supplementation in the medium, the TcPEPCK-sKO epimastigotes showed a 51% increase in ATP content compared to the tdTomato parasites. When ATP synthase in the mitochondria was inhibited in the absence of glucose, the ATP content of TcPEPCK-sKO epimastigotes was reduced by 49%. Interestingly, glucose addition, both in the presence and absence of oligomycin, resulted in similar ATP production, suggesting that, if glucose is present, ATP production in TcPEPCK-sKO epimastigotes is less dependent on mitochondria (Figure 2F).

### PEPCK deficiency increased PPDK expression and alanine production in *T. cruzi* epimastigotes

Another enzyme that competes for PEP with PEPCK in the glycosomes is the pyruvate phosphate dikinase (PPDK). This enzyme converts PEP into pyruvate, thereby promoting ATP generation from AMP and PPi. Pyruvate is then reduced to alanine by alanine dehydrogenase, regenerating one NAD^+^ inside the glycosomes. Alanine produced in this reaction can be secreted into the medium (Figure 3A). Here, PPDK expression was quantified by PCR and showed a 2-fold upregulation in TcPEPCK-sKO parasites compared to tdTomato parasites (Figure 3B). To analyze the activity of this route, intracellular and secreted alanine levels were measured in both parasite lines. Intracellular alanine increased by 60% in TcPEPCK-sKO parasites (Figure 3C). However, PEPCK deficiency led to 35% reduction in alanine secretion to the supernatant (Figure 3D). These results strongly suggest that TcPEPCK-sKO epimastigotes generate more alanine via the PPDK route, which can be used as a carbon source for the parasite, leading to reduced secretion into the supernatant.

**Figure 3.**
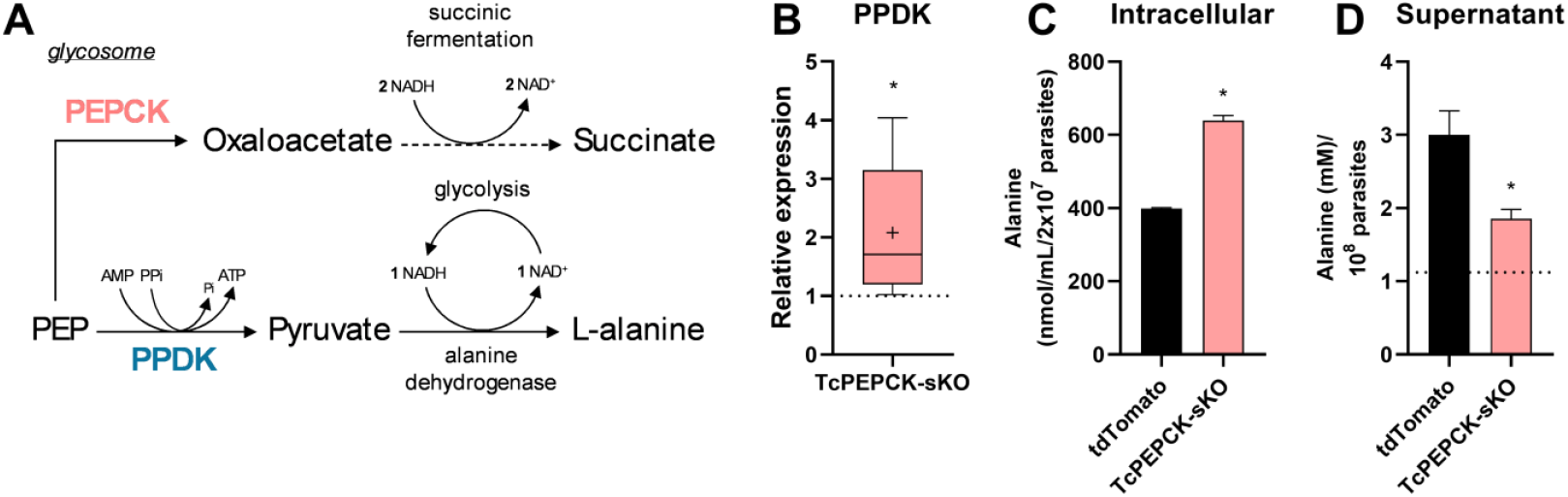
Evaluation of PPDK enzyme in TcPEPCK-sKO epimastigotes. **(A)** Pyruvate phosphate dikinase (PPDK) and PEPCK compete for the same substrate inside the glycosomes. This enzyme converts this molecule into pyruvate using inorganic pyrophosphate (PPi) and AMP to generate ATP and inorganic phosphate (Pi). Pyruvate then receives an amino group and is reduced to alanine by alanine dehydrogenase, which is then excreted. **(B)** tdTomato and TcPEPCK-sKO epimastigotes were grown in LIT medium for 7 days at 28 ºC. PPDK expression in the parasites was measured using qPCR. All quantitative measurements were performed in triplicate and normalized to the internal control TCZ (195-bp repeated DNA) for each reaction. The data are the mean ± SEM of five independent experiments analyzed by Mann-Whitney test. **(C)** Intracellular alanine levels were determined in the supernatants of frozen-thawed parasites. Representative data present mean ± SD of three independent experiments prepared in duplicates. **(D)** Alanine secretion was measured in the culture medium supernatant of tdTomato and TcPEPCK- sKO epimastigotes and normalized to parasite number. The data are the mean ± SEM of four independent experiments. *p<0.05, compared to tdTomato, analyzed using Student’s t- test.

### Disruption of the PEPCK gene impaired the metacyclogenesis but improved the infection of *T. cruzi* in vitro and in vivo

The capacity of TcPEPCK-sKO epimastigotes to differentiate into metacyclic trypomastigotes was impaired, presenting a 62% reduction compared to tdTomato parasites, suggesting a role for PEPCK in metacyclogenesis (Figure 4A).

**Figure 4.**
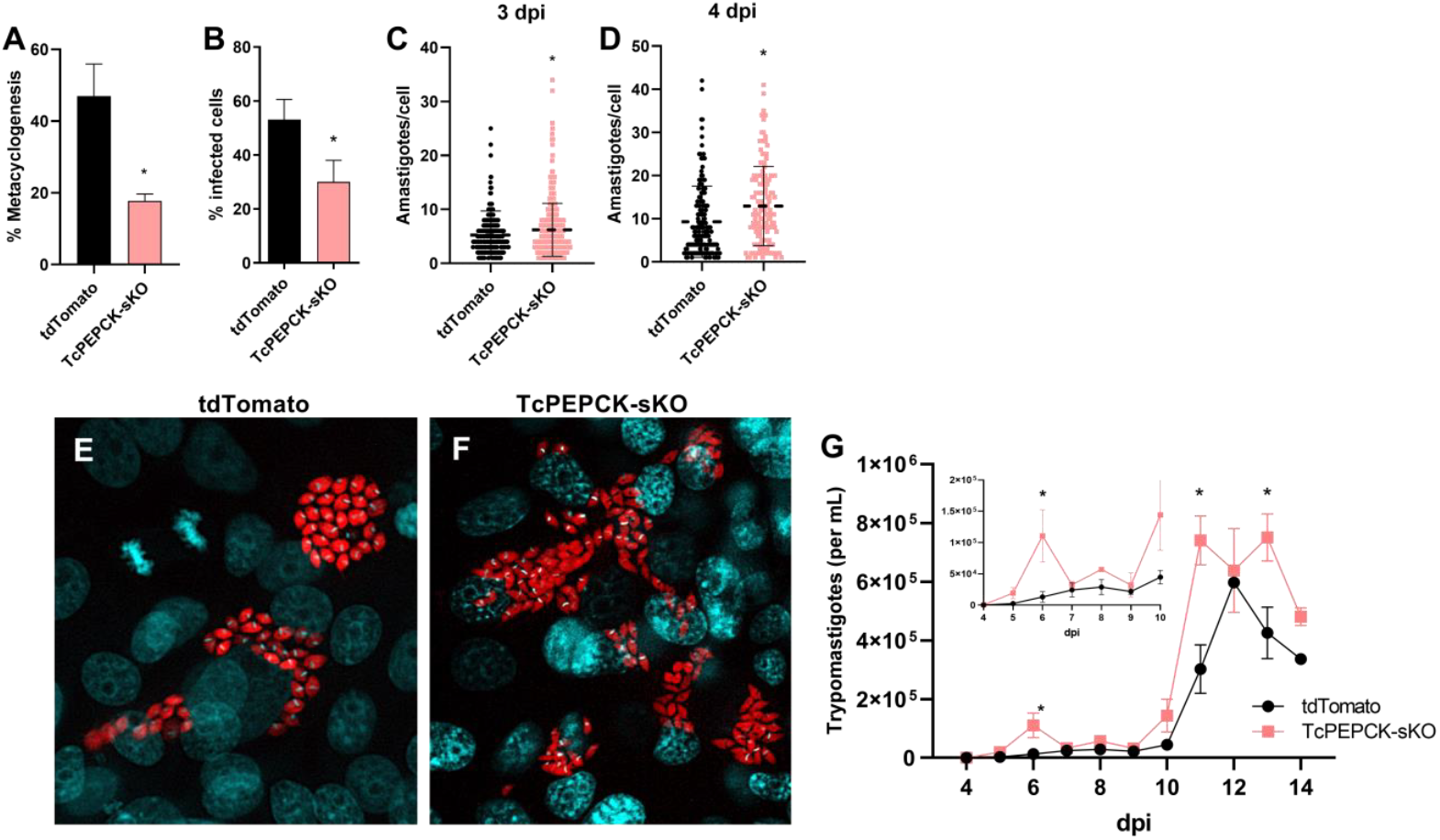
Metacyclogenesis and *in vitro* infection by TcPEPCK-sKO trypomastigotes. **(A)** For *in vitro* metacyclogenesis, tdTomato and TcPEPCK-sKO epimastigotes were challenged with TAU medium for 2 h and then diluted (1:100) in TAU3AAG medium for 96 h. The metacyclic trypomastigotes were differentiated based on their morphology and motility using a light microscope. The data are mean ± SEM of at least four independent experiments. **(B)** Vero cells were infected with tdTomato and TcPEPCK-sKO trypomastigotes for 4 h, fixed, and stained with 1 μg/mL DAPI. A total of 300 random cells were counted under a fluorescence microscope and the results were expressed as the percentage of invasion. *p<0.05, compared to tdTomato, analyzed using Student’s t-test. Amastigote replication: After 4 h of interaction and removal of trypomastigotes, new RPMI medium supplemented with 2% FBS was added, and the infected cells were maintained for **(C)** 3 dpi and **(D)** 4 dpi. The number of amastigotes per cell was counted under a fluorescence microscope by DAPI (nuclei and kinetoplast DNA) and tdTomato (amastigote cytosol) fluorescence. The data are mean ± SD from each infected cell counted in three independent experiments. *p<0.05, compared to tdTomato, analyzed using the Mann-Whitney test. Representative images show host cells infected with **(E)** tdTomato and **(F)** TcPEPCK-sKO parasites expressing tdTomato fluorescence (red) and stained with 4 dpi DAPI (blue). **(G)** Production of trypomastigotes: Trypomastigotes released into the supernatant of infected Vero cells were counted daily until 14 dpi using a Neubauer chamber. The data are mean ± SEM of at least three independent experiments. *p<0.05, compared to tdTomato parasite analyzed by multiple t-test with post-test Sidak-Bonferroni method.

Here, we demonstrated that PEPCK impairment affected the invasion of Vero cells by trypomastigotes, reducing by 43% their invasion capacity compared to tdTomato trypomastigotes (Figure 4B). Surprisingly, once inside the mammalian cells, the number of TcPEPCK-sKO amastigotes per cell increased by 19% at 3 days post-infection (dpi) (Figure 4C) and reached 43% more amastigotes at 4 dpi (Figure 4D-4F). In addition to the increase in the number of amastigotes, we also observed advanced differentiation into infective trypomastigotes in TcPEPCK-sKO parasites inside the host cell (Figure 4F) compared to tdTomato parasites (Figure 4E). We evaluated the effects of PEPCK depletion on the intracellular cycle of *T. cruzi*. TcPEPCK-sKO showed an 8-fold increase in the number of trypomastigotes released into the culture supernatant at 6 dpi (Figure 4G insert), followed by a second trypomastigote release peak at 11 dpi (Figure 4G).

To verify whether PEPCK mutant parasites were able to establish *in vivo*, IFNγ deficient mice (IFNγ-KO) were infected intraperitoneally with trypomastigotes. Similar to the *in vitro* results, the parasitemia of the IFNγ-KO mice infected with TcPEPCK-sKO trypomastigotes was much higher than the parasitemia of those infected with tdTomato parasites (Figure 5A). In animals infected with TcPEPCK-sKO, an increase of 8.9-fold and 13.6-fold in the number of trypomastigotes in the bloodstream was observed at 7 and 10 dpi, respectively, compared to tdTomato-infected mice. At 12 dpi, parasitemia decreased and again increased 8.6-fold at 15 dpi (Figure 5A). At 16 dpi, the animals were sacrificed to evaluate the tissue infection. The number of nests and parasitic loads in the heart, skeletal muscle, and gut were determined by imaging the clarified tissues and qPCR. Representative images of the clarified tissues allowed us to observe different numbers of nests in each tissue (Figure 5B). At 16 dpi, the number of nests on the skeletal muscle was higher than that on the other tissues in both tdTomato and TcPEPCK-sKO mice. However, the number of nests in TcPEPCK-sKO-infected tissues was much lower than in tdTomato- infected tissues (Figure 5B, 5C). Parasite loads measured by qPCR confirmed these findings (Figure 5D). Therefore, despite the high parasitemia in animals infected with PEPCK-mutant parasites (Figure 5A), the tissues of these animals presented lower parasite levels than those of tdTomato-infected animals (Figure 5C and 5D).

**Figure 5.**
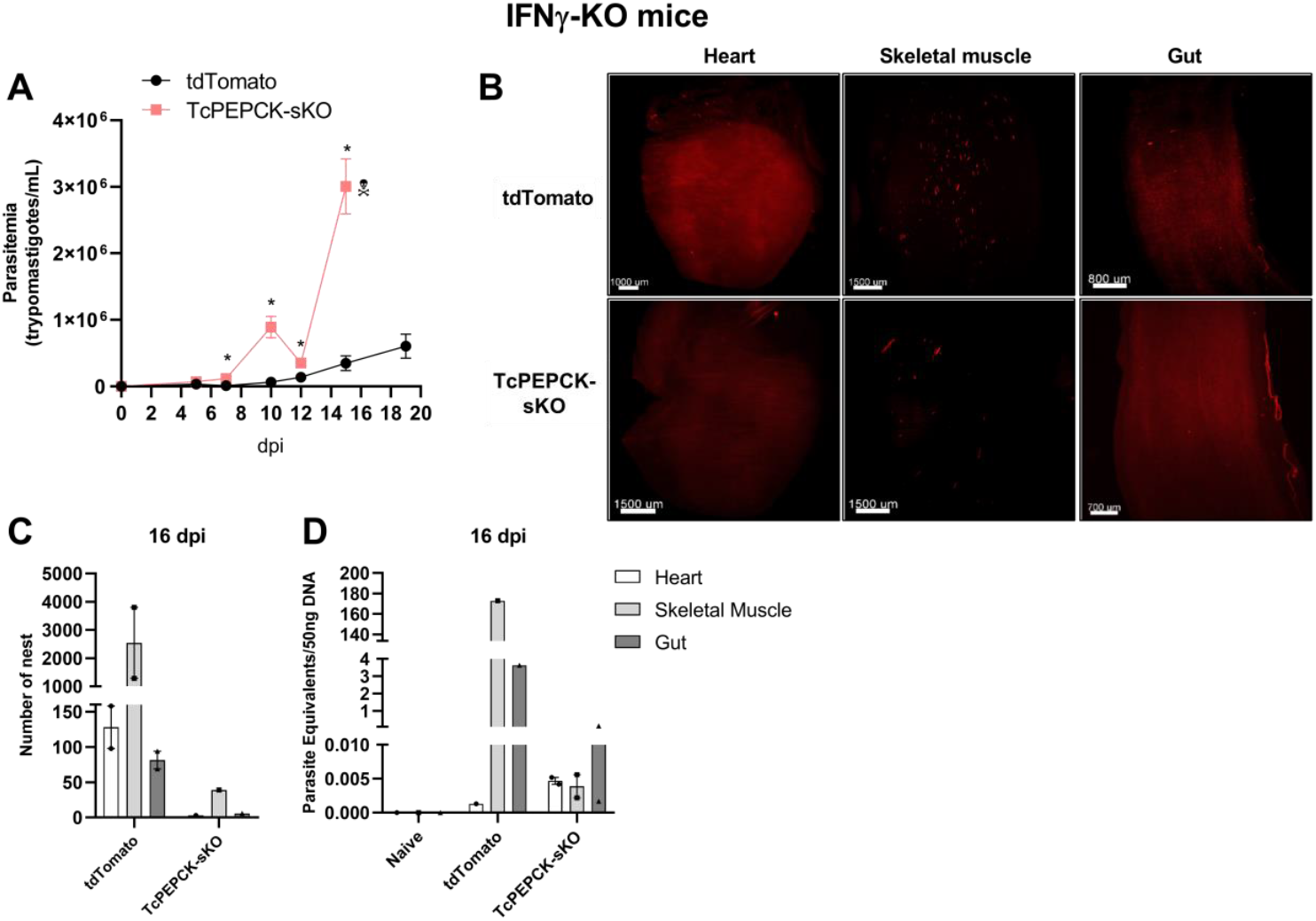
*In vivo* infection of immunodeficient mice by TcPEPCK-sKO trypomastigotes. C57BL/6 IFNγ-KO mice were infected intraperitoneally with 2 × 10^5^ tdTomato or TcPEPCK-sKO trypomastigotes. **(A)** Parasitemia was monitored from 5 to 15 days post infection (dpi). Results are presented as the mean ± SEM of parasitemia in four mice. *p<0.05, compared to tdTomato-infected mice analyzed by multiple t-tests with post-test Sidak-Bonferroni method. **(B)** Heart, skeletal muscle, and gut samples from mice infected with tdTomato or TcPEPCK-sKO were obtained after 16 dpi, clarified using the CUBIC protocol, and imaged using LSFM. **(C)** Automated quantification of total *T. cruzi* amastigote nests in 3D reconstructed tissues using the tdTomato fluorescent protein expressed in parasites. **(D)** Parasitic loads in tissues were determined by qPCR after 16 dpi.

We also evaluated *in vivo* infection of immunocompetent mice. In these animals, the presence of trypomastigotes in the blood was observed at 5 and 7 dpi in tdTomato and TcPEPCK-sKO parasites; however, there was no significant difference in parasitemia (Figure 6A). At 10 dpi, trypomastigotes were not detected in the bloodstream. The infected mice were sacrificed at 21 dpi for tissue analysis. We observed no significant changes in the number of parasite nests (Figure 6C) or the parasite load inside the tissues using LSMS (Figure 6B) or qPCR (Figure 6D). In the heart and skeletal muscles, the numbers of tdTomato and TcPEPCK-sKO parasites were the same (Figure 6D). Although the TcPEPCK-sKO parasite is also capable of establishing infection in wild-type animals, the active immune system efficiently controls parasite growth, and its dissemination is observed in immunodeficient animals (Figure 5).

**Figure 6.**
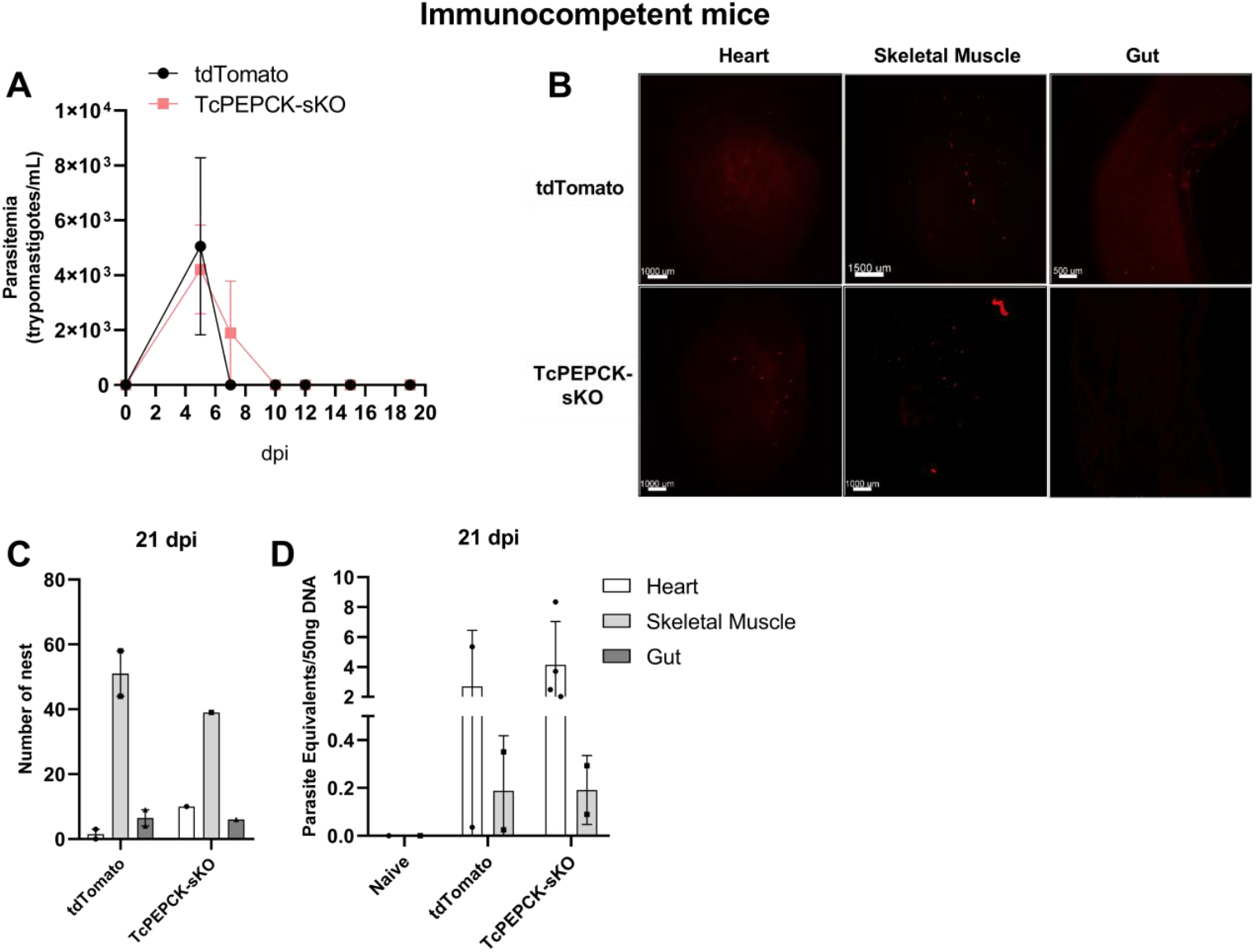
*In vivo* infection of immunocompetent mice by TcPEPCK-sKO trypomastigotes. Wild-type C57BL/6 mice were intraperitoneally infected with 2 × 10^5^ tdTomato or TcPEPCK-sKO trypomastigotes. **(A)** Parasitemia was monitored from 5 to 21 days post infection (dpi). Results are presented as the mean ± SEM of parasitemia in four mice. *p<0.05, compared to tdTomato-infected mice analyzed by multiple t-tests with post-test Sidak-Bonferroni method. **(B)** Heart, skeletal muscle, and gut samples from mice infected with tdTomato and TcPEPCK-sKO were obtained after 21 dpi, clarified using the CUBIC protocol, and imaged using LSFM. **(C)** Automated quantification of total *T. cruzi* amastigote nests in 3D reconstructed tissues using the tdTomato fluorescent protein expressed in parasites. **(D)** Parasitic loads in tissues were determined using qPCR at 21 dpi.

## Discussion

The importance of PEPCK as an essential gene has been previously proposed based on genome-scale metabolic models **[18]**. Here, deletion of the glycosomal PEPCK gene in *T. cruzi*, mediated by the CRISPR/Cas9 system, resulted in clones containing only one allele of the gene. This observation suggests that complete loss of the PEPCK gene might render non-viable parasites, indicating that this could be an essential gene for the epimastigote stage of *T. cruzi* when grown in regular medium. Partial removal of the gene led to a decrease in PEPCK activity, which catalyzes the carboxylation of PEP to oxaloacetate, generating ATP in the glycosomes of the parasite, and potentially affecting the subsequent reactions of the succinic fermentation branch in glycosomes **[12]**. In the procyclic form of *T. brucei*, succinic fermentation is the main pathway used to maintain the redox and ATP/ADP balance in the glycosomes, and this function is likely to extend to *T. cruzi* as well **[19,20]**. Succinate, the most excreted molecule produced by glucose metabolism in *T. cruzi*, is produced by fermentation **[12,20]**. Our results showed that PEPCK deficiency reduced glucose uptake by epimastigotes, indicating that this impairment perturbed glycolytic flux. It has been shown that inhibition of PEPCK (through noncompetitive inhibition or gene knockout) also results in decreased glucose consumption and succinate production and secretion, with an accumulation of alanine in *T. cruzi* **[20]**, and *T. brucei*. Interestingly, these metabolic changes did not alter *T. cruzi* epimastigote growth in our study, and the same phenomenon was observed in the procyclic forms of *T. brucei* **[19]**.

With the disruption of PEPCK, *T. cruzi* has alternative routes to NADH reoxidation inside the glycosome, one of which is initiated by PPDK. In our study, PEPCK impairment promoted an increase in PPDK expression, resulting in increased levels of alanine inside the cells. In non-deleted parasites, the produced alanine is usually excreted into the environment **[16,21]**, but here, the ablation of PEPCK caused a reduction in excretion, possibly indicating that the parasite can use alanine as a carbon source. A similar result was observed by Urbina et al. **[20]**, who reported that alanine generation per mole of glucose consumed increased when PEPCK was inhibited. The PEPCK-related pathway reoxidizes two NADH molecules, whereas PPDK reoxidizes one. These diminished NADH-consuming reactions can influence glycolytic activity, and consequently, glucose uptake by the parasite, as observed in the present study.

Oxidative phosphorylation in the mitochondria is another mechanism of energy acquisition in organisms. Our data demonstrated that partial deletion of PEPCK led to decreased glucose consumption (36%) and impaired oxidative phosphorylation (43% decrease). However, total ATP production was not significantly altered even in the presence of the mitochondrial ATP synthase inhibitor, oligomycin. This could explain why TcPEPCK-sKO parasites seemed to be less dependent on ATP synthesis through mitochondrial respiration when glucose was supplied as a fuel, as shown in Figure 2. In addition to glucose, trypanosomes use amino acids to meet their energy needs, and proline is one of their preferred amino acids **[15,22]**. When PEPCK is inhibited, proline catabolism is significantly affected by *T. cruzi* **[20]**. In *T. brucei* procyclic form, otherwise, when they are PEPCK knocked out or glucose-depleted medium, the parasite switches its metabolism and increases proline consumption to sustain a portion of energy generation through mitochondrial oxidation pathways **[19]**. Oxaloacetate, which is necessary for PEPCK- driven gluconeogenesis (generation of hexose phosphates) and likely derived from the carbon skeleton of proline, is affected because pyruvate from glucose is required **[20]**. This affects the rate of production of TCA cycle metabolites, possibly causing a reduction in the mitochondrial respiration of the parasite, as shown in the present study.

In the life cycle of *T. cruzi*, epimastigotes differentiate into infective metacyclic trypomastigotes under nutritional stress. Here, PEPCK was shown to be crucial for metacyclogenesis as it reduced the differentiation of TcPEPCK-sKO parasites by 62%. During metacyclogenesis, the expression of genes related to energy acquisition via glucose oxidation is downregulated **[23]**. However, intracellular succinate levels increase over time during differentiation **[24]**. With reduced glucose metabolism, succinate can be generated in the TCA cycle in mitochondria, fueled by proline and glutamine oxidation, both of which are known to stimulate *T. cruzi* metacyclogenesis **[25,26]** or by succinic fermentation in glycosomes, eventually when metabolites from amino acid catabolism are driven to gluconeogenesis. PPDK and PEPCK are key enzymes for gluconeogenesis in *T. brucei*, and the inhibition of this anabolic pathway is essential for the differentiation of infective metacyclic forms in tsetse flies **[27]**.

Proteomic analysis of *T. cruzi* amastigotes showed that PEPCK is expressed during this replicative stage **[28]**. Surprisingly, intracellular amastigotes with compromised PEPCK activity show significantly increased replication, driving faster differentiation into trypomastigotes. This enhanced infection resulted in a higher production of trypomastigotes in the supernatant, suggesting an improvement in the virulence of TcPEPCK-sKO. Similarly, *in vivo* experiments showed that TcPEPCK-sKO trypomastigotes were easily established in immunodeficient IFNγ-KO mice, causing high parasitemia and rapid debilitation. Notably, although there were high levels of parasites in the mouse bloodstream, the parasitic loads inside the tissues were lower than those in the tissues infected with tdTomato parasites. This result indicated that TcPEPCK-sKO parasites presented faster and enhanced intracellular cycles inside the tissues, leading to higher trypomastigote release into the bloodstream, which could not be measured on the last day of the experiment in these samples. Padilla et al. **[29]** showed that when infection occurs in the footpad, the skeletal muscle is one of the first tissues infected by *T. cruzi* followed by a gradual decline accompanied by an increase in parasite load in the heart of the animals. Therefore, the low parasite loads observed inside the tissues may represent the peak infection profile, with animals infected with TcPEPCK-sKO parasites reaching their peak infection earlier than those infected with tdTomato parasites. Conversely, in immunocompetent mice, the immune system effectively controlled *T. cruzi* growth, with a peak infection at 5 dpi, similar to that in tdTomato-infected mice.

Contrary findings were observed in *Leishmania sp*. infection in mammalian cells; the absence of PEPCK dramatically impaired the replication of *L. major* amastigotes inside macrophages and promoted attenuation both *in vitro* and *in vivo*, resulting in no stimulation of the immune system, as observed when PEPCK was naturally expressed in the parasites **[30]**. *Leishmania* species use enzymes such as PPDK, PEPCK, and glycerol kinase (GK) to synthesize carbohydrates from non-carbohydrate molecules to survive in sugar-starved environments, such as the phagolysosomes of macrophages **[31]**.

Although *Leishmania* spp. and *T. cruzi* are both trypanosomatids, these protozoa employ distinct metabolic pathways and roles that may vary in their survival, differentiation, or infectivity depending on specific and restrictive environmental conditions. Unlike *Leishmania, T. cruzi* amastigotes proliferate in the cytosol of host cells and rely on available nutrients there **[32]**. In response to nutrient fluctuations, this intracellular stage modifies a subset of metabolic genes and the host cell metabolism to replicate inside the cell **[33,34]**. The primary carbon sources for amastigote energy metabolism may be fatty and amino acids complemented by glucose metabolism. Fatty acid oxidation, the TCA cycle, oxidative phosphorylation-related genes and proteins, and amino acid catabolism are upregulated at the intracellular stage during development **[28,35]**. However, glycolysis genes were initially reduced and then upregulated after 24 h of amastigote development compared to trypomastigotes **[35]**. The need for glucose in amastigotes induces the modulation of mammalian cell metabolism, increasing glucose consumption and metabolism over time. This parasite scavenges glucose for ATP production and other anabolic pathways **[34]**. Interestingly, enhanced glycolysis gene expression and activity were not accompanied by changes in fermentation-related gene expression, including that of PEPCK **[35]**. This suggests that, in addition to succinic fermentation, other pathways involving NADH-consuming reactions are active in the glycosomes of *T. cruzi* amastigotes. Since the PEPCK gene could not be completely knocked out in *T. cruzi*, alternative routes may compensate for the redox balance of the organelle. Furthermore, the partial impairment of PEPCK may increase the necessity of mitochondrial oxidative phosphorylation at the intracellular mammalian stage, thereby enhancing amastigote replication and *T. cruzi* virulence.

## Conclusion

Figure 7 presents the proposed scheme of metabolic changes in *T. cruzi* epimastigotes with impaired PEPCK activity and their effects throughout the protozoan life cycle. The reduction in PEPCK activity, mediated by gene editing, compromised glucose consumption and mitochondrial physiology in epimastigotes, stimulating the use of an alternative fermentation pathway to maintain glycosomal redox status. These metabolic changes did not interfere with ATP generation, thereby allowing epimastigote proliferation. However, metacyclogenesis was diminished by disruption of PEPCK. TcPEPCK-sKO trypomastigotes showed difficulty invading mammalian cells, but this effect was overcome upon differentiation into amastigotes. The replication of PEPCK-impaired amastigotes increased over time, resulting in increased generation of trypomastigotes *in vitro*. This effect has also been observed in animals, where a single PEPCK knockout led to uncontrolled parasitemia and pathology in immunodeficient animals. In summary, PEPCK appears to play a central role in the bioenergetics of *T. cruzi* epimastigotes and other life stages by regulating essential biological processes such as metacyclogenesis and infection capacity.

**Figure 7.**
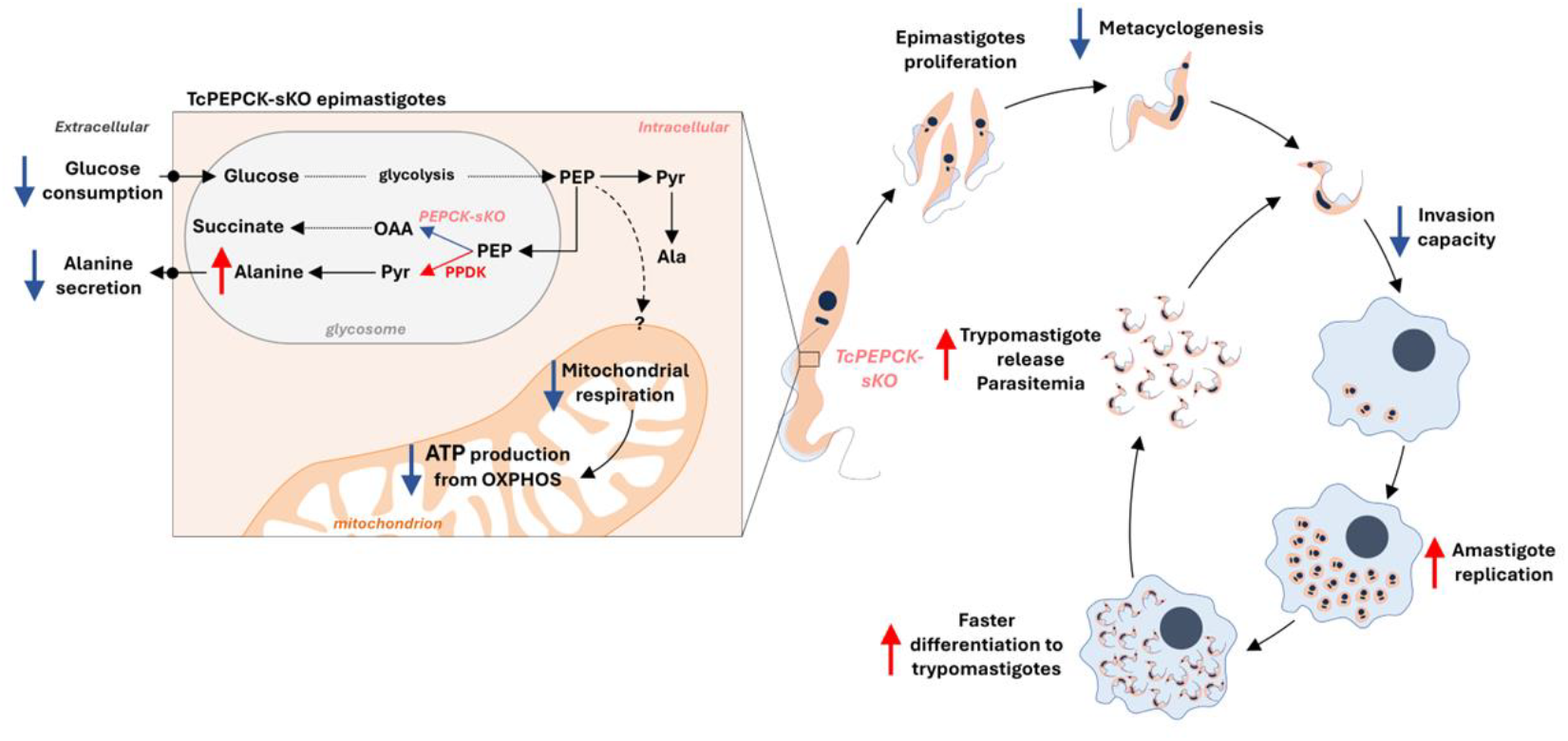
Schematic representation of the effects of PEPCK deficiency in *T. cruzi* bioenergetics and life stages. CRISPR/Cas9 editing resulted in a single-allele *T. cruzi* mutant of PEPCK. 1) In epimastigotes, partial gene removal resulted in decreased PEPCK activity (and probably succinic fermentation), which reduced glycolytic flux and mitochondrial respiration when glucose is available. In response, the PPDK fermentation route was stimulated, leading to enhanced production of intracellular alanine, which is mainly retained inside the cell. 2) These metabolic perturbations adapt the epimastigote form to supply sufficient energy for proliferation, but not to differentiate into metacyclic trypomastigotes. 3) PEPCK impairment also reduced the invasive capacity of the infective stage. However, once inside the host cell, *T. cruzi* amastigotes replicate faster when PEPCK-mediated fermentation is affected, accelerating the differentiation and production of infective-stage trypomastigotes. Infection with PEPCK-deficient parasites induced higher parasitemia and dissemination in immunodeficient animals; however, this was controlled by an active immune system. Overall, the PEPCK enzyme was shown to be important in the regulation of *T. cruzi* energy metabolism, reducing the differentiation and invasion of infective forms, but improving infection by boosting amastigote replication.

## Methodology

### Mammalian cells and parasite culture

Vero cells were cultured in RPMI 1640 medium (Vitrocell) (containing 4,7 g/mL HEPES, 0.2% NaHCO3 (w/v), 200 units/mL penicillin, 200 μg/ml of streptomycin, 25 μg/mL gentamicin, 1 mM sodium pyruvate and 50 μM 2-βmercaptoethanol) supplemented with 10% FBS in a humid atmosphere containing 5% CO_2_, at 37 °C .

Epimastigote forms of *T. cruzi* (Brazil A4 strain) expressing tdTomato were obtained as previously described **[8]** and maintained at 28 °C in liver infusion tryptose (LIT) medium, supplemented with 10% fetal bovine serum (FBS), 30 μM heme and 250 μg/ml G418.

Culture-derived trypomastigotes were obtained from infected Vero cells, as previously described by Canavaci et al. **[36]**. Briefly, the cells were infected with tdTomato or TcPEPCK-sKO metacyclic trypomastigotes obtained through *in vitro* metacyclogenesis previously purified in non-inactivated FBS at 37 ºC. After 24-48 h of interaction, the infected cells were gently washed with PBS to remove non-internalized trypomastigotes and maintained in RPMI supplemented with 10% FBS at 37 °C in 5% CO_2_. Trypomastigotes were collected 5 dpi and used in subsequent experiments.

### sgRNA preparation

The sgRNA target sequences and corresponding repair templates were identified using the Eukaryotic Pathogen CRISPR guide RNA/DNA Design Tool (http://grna.ctegd.uga.edu) with the SaCas9 option (21-bp target sequence preceding an NNGRRT PAM site) (Table S1). sgRNAs were prepared as previously described **[8]**. Briefly, DNA templates for sgRNA *in vitro* transcription (IVT) were generated by PCR to amplify the sgRNA scaffold sequence specific for SaCas9 using the TranscriptAid T7 high-yield transcription kit (Thermo Scientific), and sgRNAs were purified by ethanol precipitation.

### SaCas9 and sgRNA assembly and electroporation

SaCas9 and sgRNAs were prepared immediately before transfection. SaCas9 expression and purification were performed as described by Soares Medeiros et al. **[10]**. The sgRNAs were annealed by heating to 90 °C for 5 min and slowly cooled to room temperature. Equimolar amounts of SaCas9 and sgRNA (1:1 ratio) were incubated at room temperature for 10 min. *T. cruzi* epimastigotes in early log phase (2.5 × 10^6^ parasites) were resuspended in 100 μl room temperature human T cell Nucleofector solution (Amaxa AG). The DNA repair template (40 ng) was added as indicated. Electroporations were performed using an Amaxa Nucleofector device; the program X-14 was used to electroporate the parasites. Electroporated *T. cruzi* epimastigotes were cultured in 25 cm^2^ cell culture flasks with 5 ml LIT medium.

### Generation of TcPEPCK gene knockout

*T. cruzi* epimastigotes of the Brazil strain in log-phase growth were transfected with RNP complexes containing sgRNAs designed to knock out PEPCK-encoding genes (TcCLB.507547.90, TcBrA4_0071790, and TcBrA4_0071840), as well as 40 ng of repair template containing three in-frame stop codons and the M13 bacteriophage sequence (Table S1). Single parasites were flow-sorted at 8 days post-transfection (d.p.t.) into individual wells of 96-well plates containing 50% LIT medium and 50% parasite-conditioned medium. DNA from transfected parasites was isolated at 15 and 30 d.p.t. from the uncloned population and at 40 days post-cloning to assess genomic integration of the repair template using PCR.

### PEPCK enzyme activity

PEPCK activity was measured following the protocol provided with the Phosphoenolpyruvate Carboxykinase Activity Assay Kit (Sigma-Aldrich). Proteins from TdTomato and TcPEPCK-sKO epimastigote of *T. cruzi* were extracted after 5 cycles of freezing/thawing in a PEPCK-specific buffer. The protein concentration was determined using a bicinchoninic acid (BCA) protein assay kit (Thermo Scientific), with bovine serum albumin as the standard. The generation of the final product was monitored using spectrophotometry for 60 min.

### Epimastigotes proliferation

Epimastigotes tdTomato or TcPEPCK-sKO (2.5 × 10^6^ parasites/mL) were grown at 28 °C for 7 days in LIT medium supplemented with 10% FBS and 30 μM heme in the presence or absence of 10 mM glucose (Sigma-Aldrich). Parasite growth was monitored by counting cells in a Neubauer chamber.

### Glucose consumption and alanine secretion

tdTomato and TcPEPCK-sKO epimastigotes (1 × 10^8^ parasites/mL) were incubated in LIT medium for 24 h. The amount of glucose in the supernatant was determined using a glucose liquid form kit (Labtest, Minas Gerais, Brazil). The alanine concentration in the supernatant was determined using an Alanine Assay Kit (Sigma-Aldrich, Germany). Glucose consumption and alanine secretion were determined by calculating the difference between the glucose or alanine concentrations in the cell-free medium and the supernatant after parasite culture. Values were normalized to the number of cells.

### Oxygen consumption rates

The O_2_ consumption rates (OCR) of tdTomato and TcPEPCK-sKO epimastigotes were assessed using high-resolution respirometry (Oxygraph 2 K; Oroboros Instruments, Innsbruck, Austria) under continuous stirring. The temperature was maintained at 28 °C and intact parasites (5 × 10^7^ parasites/chamber) were added in 2 mL of respiration buffer (125 mM sucrose, 65 mM KCl, 2 mM KH_2_PO_4_, 0.5 mM MgCl_2_, 10 mM HEPES-KOH, and 1 mg/mL fat acid-free albumin; pH 7.2) **[37]**. The rate of O_2_ consumption by parasites in their natural physiological state (basal) was measured for at least 10 min. Leak respiration was reached after the addition of 2 μg/mL oligomycin. Uncoupled state of maximum respiration was stimulated by the addition of increasing concentrations of carbonyl cyanide-p-(trifluoromethoxy) phenylhydrazone (FCCP), up to 3.5 μM (Sigma-Aldrich, St. Louis, MO, USA) to allow the maximal capacity of the electron transport system (ETS) to be measured. ETS-related respiration was inhibited by the addition of 3 μg/mL Antimycin A (AA), a complex III inhibitor (Sigma-Aldrich, St. Louis, MO, USA) to determine residual oxygen consumption (ROX). The physiological mitochondrial OCR (routine), leak respiration, and ETS maximal capacity were calculated by subtracting the ROX consumption values from the initial OCR and after the addition of oligomycin and FCCP, respectively. Oligomycin blocks the F_0_ subunit of mitochondrial ATP synthase, allowing the calculation of OXPHOS-related OCR (the difference between Routine and Leak respiration) **[38]**.

### ATP quantification

tdTomato or TcPEPCK-sKO epimastigotes were submitted to LIT medium without the supplementation of glucose for 24 h. Then, the parasites were challenged with or without 10 mM glucose or 2 μg/mL oligomycin for 24 h. After the incubation, the parasites were washed in PBS, and the parasite concentration was adjusted to 1 × 10^7^ cells in 400 μL of PBS. Then, we transferred 100 μL (2.5 × 10^6^ epimastigotes) onto opaque-walled 96-well plates, and the intracellular ATP was measured using ATPlite™ Luminescence ATP Detection Assay System (PerkinElmer). Luminescence was measured using a BioTek Synergy Hybrid Multi-Mode reader (BioTek) and ATP concentrations were calculated using an ATP standard curve.

### qPCR for gene expression

The expression of pyruvate phosphate dikinase (PPDK) was evaluated by quantitative real-time PCR (qPCR). *T. cruzi* epimastigotes (tdTomato or TcPEPCK-sKO) were maintained in LIT medium for 7 days at 28 ºC. Total RNA was extracted from the samples using TRIzol reagent (Invitrogen, USA). cDNA synthesis was performed using High-Capacity cDNA Reverse Transcription Kit (Applied Biosystems, USA) using 2 μg of RNA. qPCR was performed using SYBR Green PCR Master Mix (QIAGEN, Hilden, Germany). cDNA samples were used in triplicate for each experimental group and assayed with at least five independent replicates. All quantitative measurements were normalized to the internal TCZ control for each reaction **[39]**. Results were expressed as mean value ± standard error (SEM). The mRNA fold change was calculated by the following equations: ΔC_T_ = ΔC_T_ (target) - ΔC_T_ (TCZ); ΔΔC_T_ = ΔC_T (TcPEPCK-sKO)_ − ΔC_T (tdTomato)_; mRNA fold change = 2^-ΔΔCT^ **[40]**.

### Intracellular alanine pool

tdTomato or TcPEPCK-sKO epimastigotes (1 × 10^8^ parasites/mL) were incubated in LIT medium at 28 ºC for 24 h. Afterward, 2 × 10^7^ parasites were washed twice in PBS, resuspended in 200 μL PBS, and subjected to five freeze-thaw cycles. The freeze-thawed parasites were centrifuged at 14,000 RPM for 5 min and the supernatants were collected for quantification. The intracellular alanine pool was determined using an Alanine Assay Kit (Sigma-Aldrich) according to the manufacturer’s instructions.

### *In vitro* metacyclogenesis

For the induction of *in vitro* metacyclogenesis (differentiation from epimastigote to metacyclic trypomastigote forms), we used the protocol by Contreras et al. **[41]** with minor modifications. tdTomato or TcPEPCK-sKO epimastigotes of *T. cruzi* were incubated in TAU medium for 2 h at 28 °C. Subsequently, the parasites were diluted 1:100 in TAU3AAG medium in 175 cm^2^ cell culture flasks. Supernatants were collected after 96 h and metacyclic trypomastigotes were differentiated by morphology and motility under a light microscope.

### Invasion into mammalian cells

To the infectivity assays, Vero cells (2 × 10^4^ cells/well) were seeded in plates containing 12 mm round coverslips and maintained in RPMI medium supplemented with 10% FBS at 37 °C, 5% CO_2_ for 24 h. To evaluate the percentage of invasion, the cells were infected with tdTomato or TcPEPCK-sKO at a ratio of 1:10 (Vero:parasite) for 4 h and afterward washed with PBS (to remove the non-internalized trypomastigotes). Then, the coverslips containing the infected Vero cells were fixed with 4% paraformaldehyde (PFA) for 15 min, stained with DAPI (1 μg/mL) for 10 min (to visualize the parasite and host cell DNA), and washed with PBS to remove the excess of dye. Finally, the coverslips were transferred to slides and mounted with ProLong Diamond anti-fade mounting solution (Thermo Fisher Scientific, Waltham, MA, USA). Infected Vero cells were counted under an inverted fluorescence microscope (ECHO Revolve Microscope) in the presence of parasitic DNA and tdTomato inside the cytosol of 300 random cells.

### Replication of intracellular amastigotes and trypomastigotes burst

To evaluate amastigote replication, infected cells were cultured in RPMI medium supplemented with 2% FBS for 3 and 4 dpi. The cells were then fixed with 4% PFA, washed with PBS, and stained with DAPI to visualize the DNA. The coverslips were transferred to slides and mounted with ProLong Diamond anti-fade mounting solution (Thermo Fisher Scientific). The number of amastigotes per cell was determined using parasite DNA/kDNA and tdTomato in the parasitic cytosol under an inverted fluorescence microscope.

The formation of trypomastigotes was monitored daily in cultures by counting the trypomastigotes released into the supernatants using a Neubauer chamber up to 14 dpi, with medium renewal every 48 h.

### *In vivo* infection

All the mice were maintained at the University of Georgia Animal Facility under specific pathogen-free conditions. This study was conducted in strict accordance with the Public Health Service Policy on the Humane Care and Use of Laboratory Animals and the Association for the Assessment and Accreditation of Laboratory Animal Care accreditation guidelines. The study protocol was approved by the Institutional Animal Care and Use Committee of the University of Georgia.

C57BL/6 wild-type or IFNγ-KO mice were infected intraperitoneally with 2 × 10^5^ tdTomato or TcPEPCK-sKO trypomastigotes obtained from infected cultures. Parasitemia was monitored in the blood individually from the 5^th^ day of infection until the animal presented with symptoms. For the parasite number, blood slides were made with 5 μL of blood collected from the tail of each animal, and the parasitemia was calculated by counting the number of mobile parasites in 50 random fields under a ×40 objective. Mice were euthanized when they began to present with symptoms. The heart, skeletal leg muscle, and intestine were frozen at -80 °C for qPCR (to check the parasitic load in tissues) or clarified for visualization of the parasite nests through the fluorescence of tdTomato-expressing parasites using light sheet fluorescence microscopy (LSFM).

### Tissue clearing and parasite nest quantification

Tissue clearings performed in this study were done following the protocol described in Bustamante et al. **[42]**. C57BL/6 wild-type and IFN-γ KO mice were euthanized by carbon dioxide inhalation. When the animals did not present any pedal reflex, they were intracardially perfused with PBS. After perfusion, the organs were dissected, washed twice with PBS and fixed with PFA in an orbital shaker at 4 ºC overnight. The following day, organs were washed twice with PBS and subjected to the CUBIC protocol for clarification **[42]**. For image acquisition, the cleared organs were immersed in immersion oil in a quartz cuvette and prepared for imaging using LSFM. Parasite nests were quantified using tdTomato fluorescence **[43]**.

### qPCR of infected tissues

Tissue samples of skeletal muscle, heart and intestine from wild-type or IFNγ-KO mice were collected and processed for quantification of *T. cruzi* DNA by qPCR **[44]**. Tissue samples were finely minced and incubated in SDS-proteinase K lysis buffer at 55 °C. DNA was extracted twice using phenol-chloroform-isoamyl alcohol, precipitated with 100% ethanol, and resuspended in nuclease-free water. PCRs products containing iQ SYBR Green Supermix (Bio-Rad, Irvine, CA, USA) and specific primers for *T. cruzi* or mouse genomic DNA were analyzed on an iCycler. *T. cruzi* equivalents were calculated as the ratio of the *T. cruzi* satellite DNA value to the amount of tumor necrosis factor alpha (TNF- α) DNA in each sample and multiplied by 50 ng of DNA, the amount of DNA added to each PCR reaction.

### Statistical analysis

Statistical analyses were conducted using the GraphPad Prism 8 software (GraphPad Software, Inc., San Diego, CA, USA). Data are presented as the mean ± standard error (SE), and all experiments were independently performed at least three times. Data were analyzed using one-way analysis ANOVA, and differences among groups were assessed using Tukey’s post-hoc test or multiple t-tests. The unpaired Student’s t-test or Mann-Whitney test was used when necessary.

