## Supplemental Table for "Glycosomal phosphoenolpyruvate carboxykinase CRISPR/Cas9-deletion and its role in *Trypanosoma cruzi* metacyclogenesis and infectivity in mammalian host"

Table S1

| Name | Objective | Sequence (5'-3') |
|---|---|---|
| guideRNA (nt 481) | driven the SaCas9 to cut site in PEPCK ORF | TTGACGTCAGGCCGGGAATCG/ACGGGT |
| repair template | DNA donor template | GCTGGTGAGTGCAAGGCGGACCCGTCGATTTAGATAGATAGTGTAAAACGACGGCCAGTCCCGGCCTGACGTCAACGACGTGCGTGGCG |
| PEPCK-Fw | primer forward to validation of PEPCK ORF | CGGACCCGTCGATTCCCGGC |
| PEPCK-Rv | primer reverse to validation of PEPCK ORF or PEPCK knockout | AGCAATGGCCTTGCTCAGCG |
| M13-Fw | primer forward to validation the knockout | TGTAAAACGACGGCCAGT |
| M13-Rv | primer reverse to validation the knockout | ACTGGCCGTCGTTTTACA |

PAM sequence (blue)
3-in frame STOP codon sequence (red)
M13 bacteriophage (green)
homology sequence for repair template (underlined)
